## Supplementary material for "Chemokine expression profile of an innate granuloma": Figure Supplements

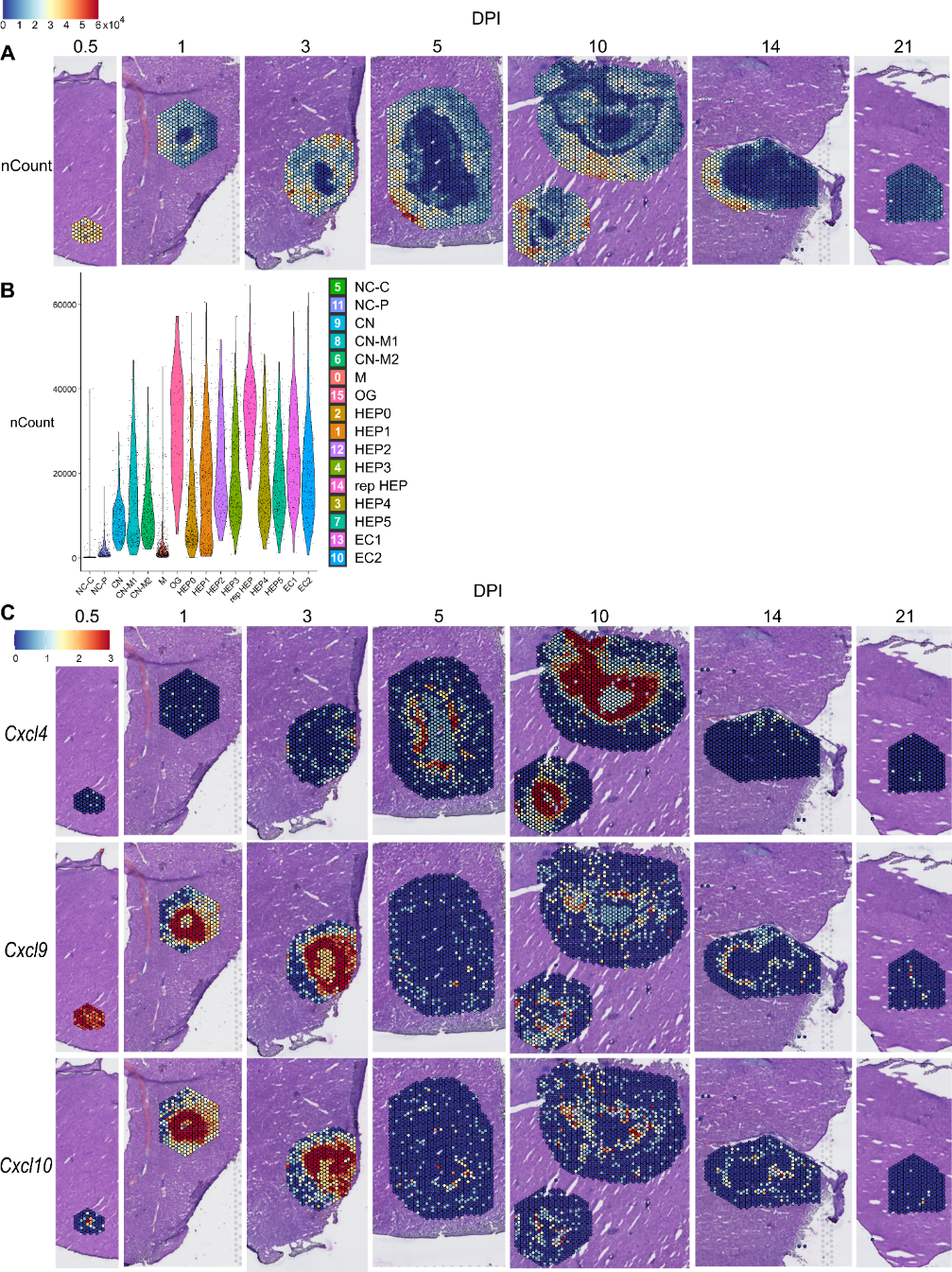

**Figure 1 – figure supplement 1. Sequencing depth of samples and spatial expression of CXCR3 ligands.** (**A**) SpatialFeaturePlot displaying raw counts (nCount) per spot at various days post-infection (DPI). Scale set at 0 – 60,000 reads. (**B**) Violin plot displaying raw counts (nCount) per cluster across all timepoints. (**C**) SpatialFeaturePlots displaying normalized gene expression data of CXCR3 ligands (i.e. *Cxcl4*, *Cxcl9*, and *Cxcl10*) at various DPI. Scale set at 0 – 3.0 expression.

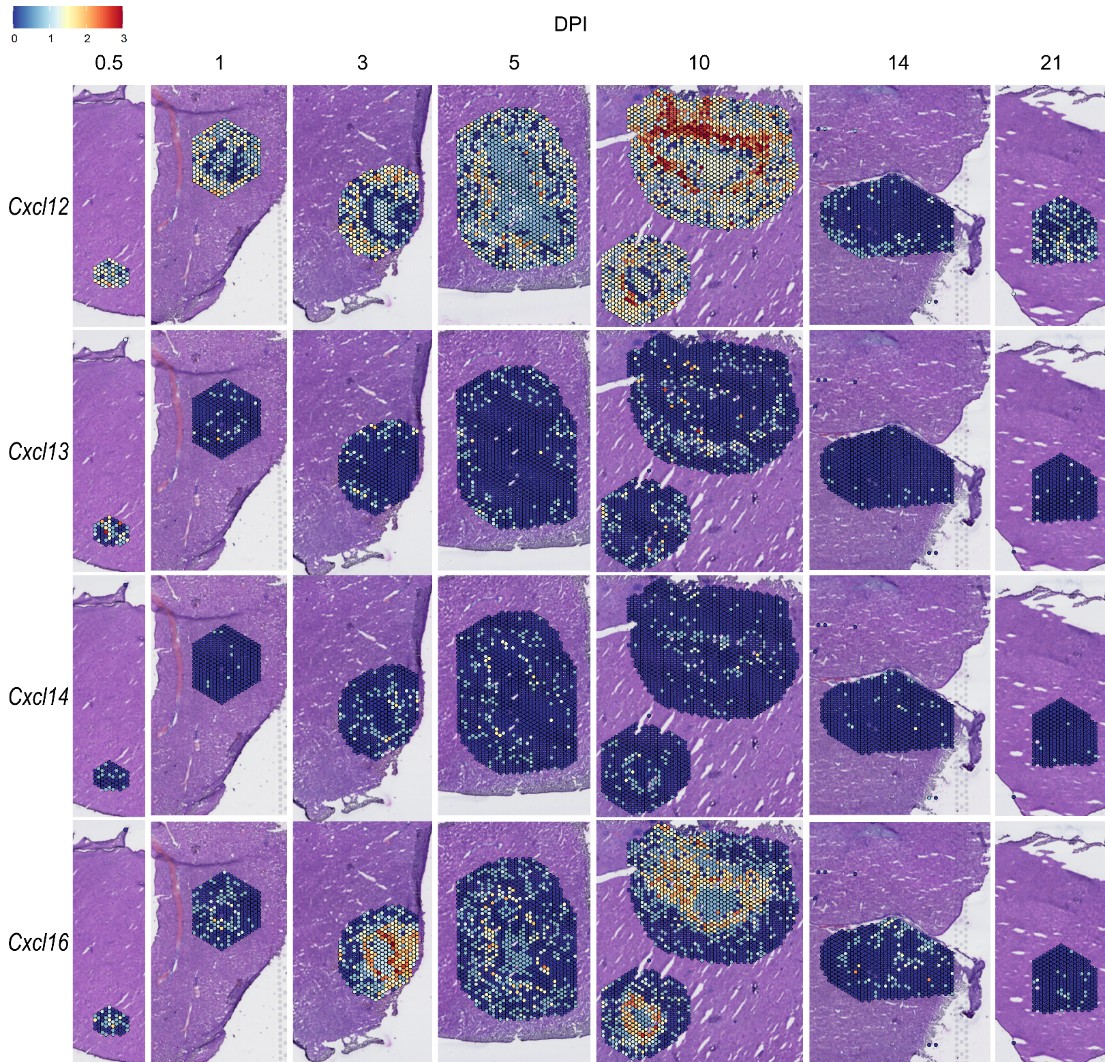

**Figure 4 – figure supplement 1. Spatial expression of *Cxcl12*, *Cxcl13*, *Cxcl14,* and *Cxcl16*.** SpatialFeaturePlots displaying normalized gene expression data of selected *Cxcl* family members at various days post-infection (DPI). Scale set at 0 – 3.0 expression.

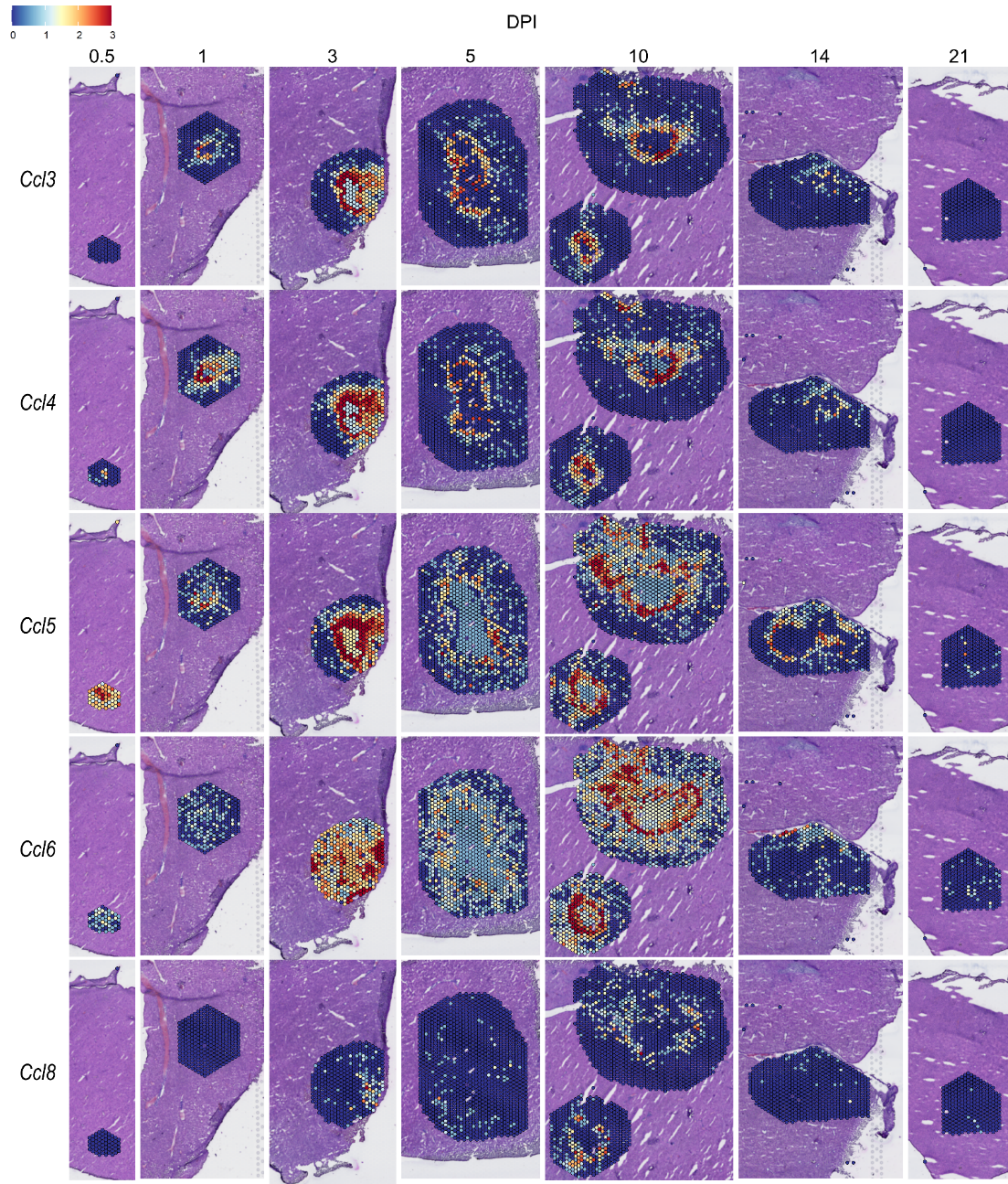

**Figure 4 – figure supplement 2. Spatial expression of *Ccl3, Ccl4, Ccl5, Ccl6,* and *Ccl8*.** SpatialFeaturePlots displaying normalized gene expression data of selected *Ccl* family members at various days post-infection (DPI). Scale set at 0 – 3.0 expression.

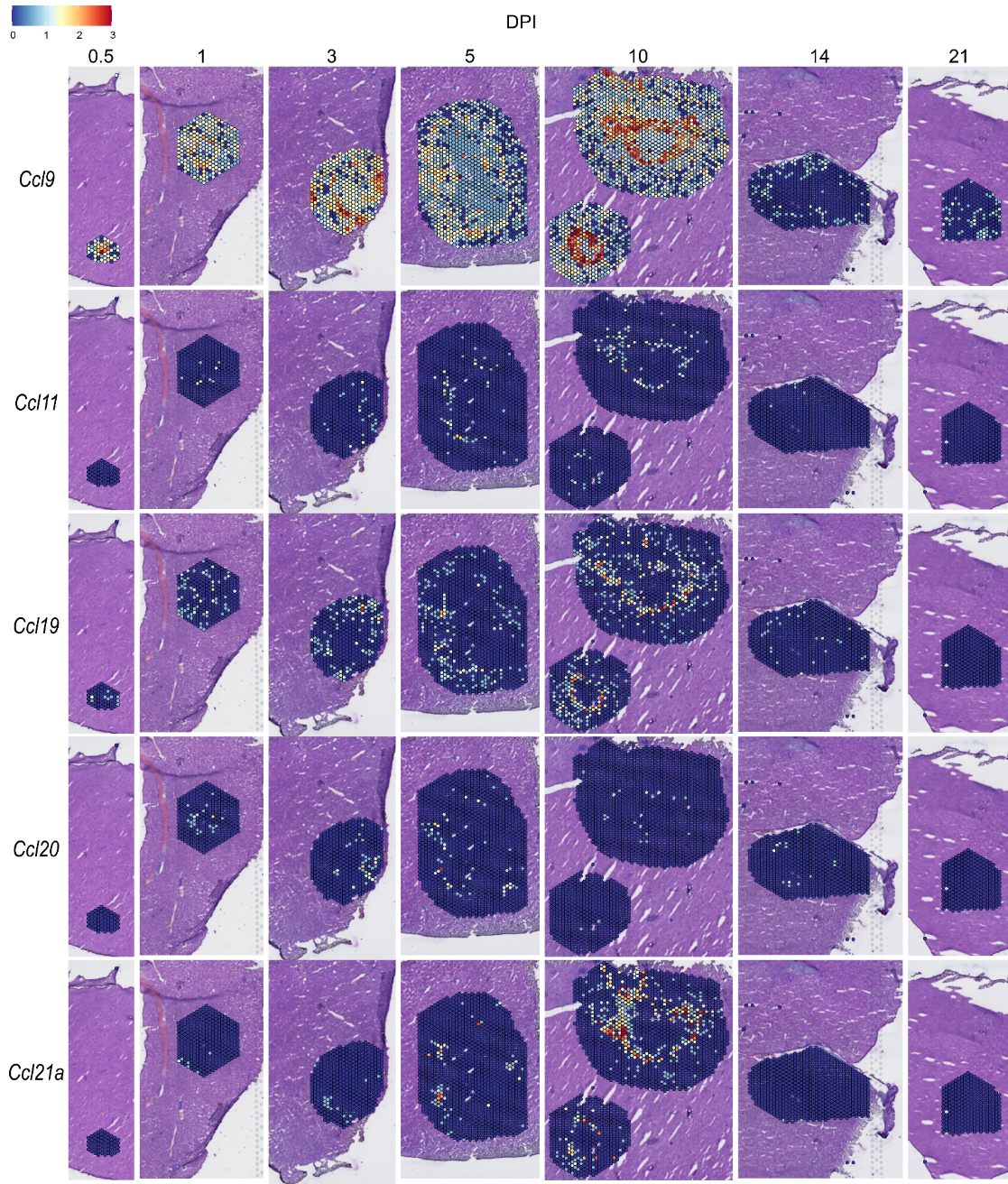

**Figure 4 – figure supplement 3. Spatial expression of *Ccl9, Ccl11, Ccl19, Ccl20,* and *Ccl21a.*** SpatialFeaturePlots displaying normalized gene expression data of selected *Ccl* family members at various days post-infection (DPI). Scale set at 0 – 3.0 expression.

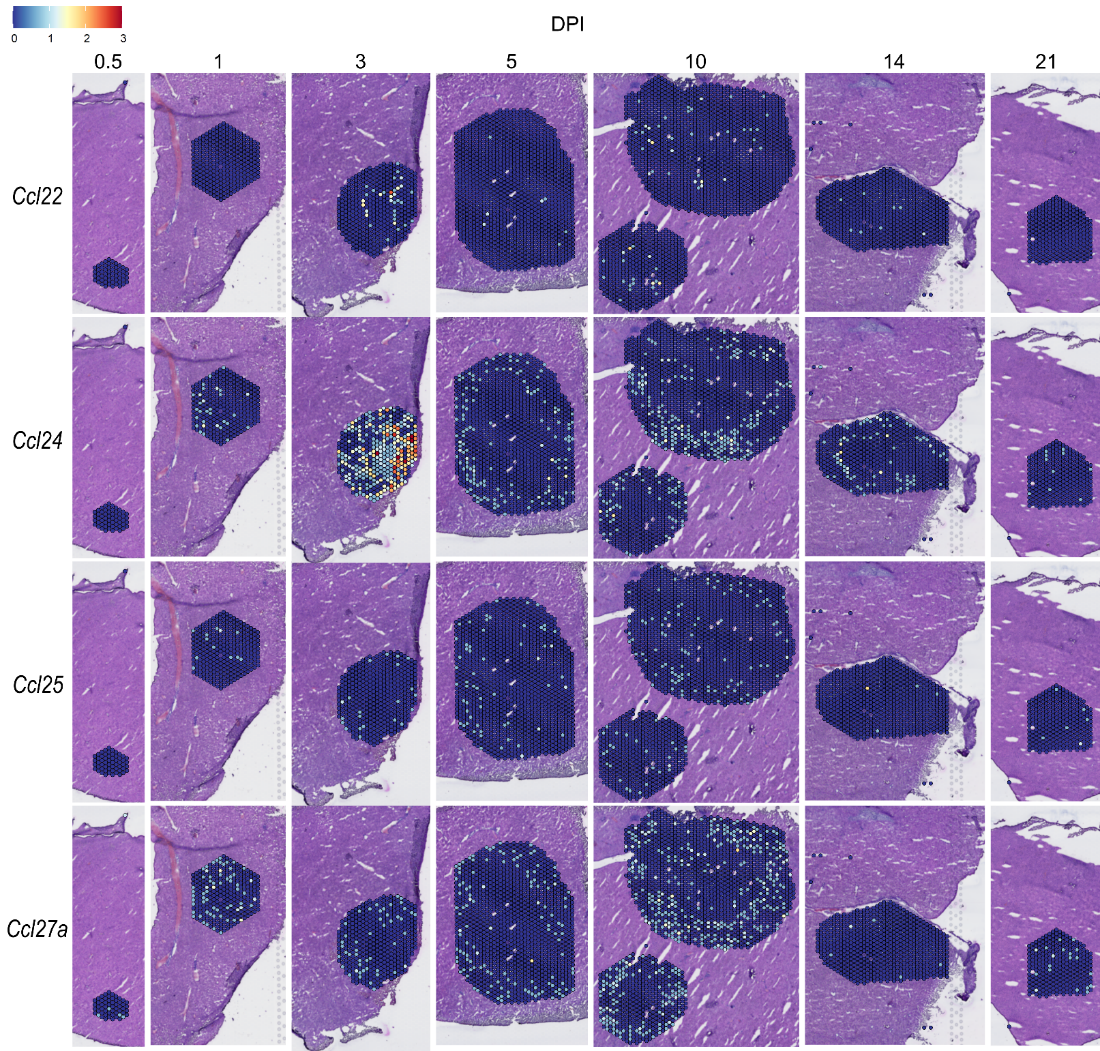

**Figure 4 – figure supplement 4. Spatial expression of *Ccl22, Ccl24, Ccl25,* and *Ccl27a*.** SpatialFeaturePlots displaying normalized gene expression data of selected *Ccl* family members at various days post-infection (DPI). Scale set at 0 – 3.0 expression.

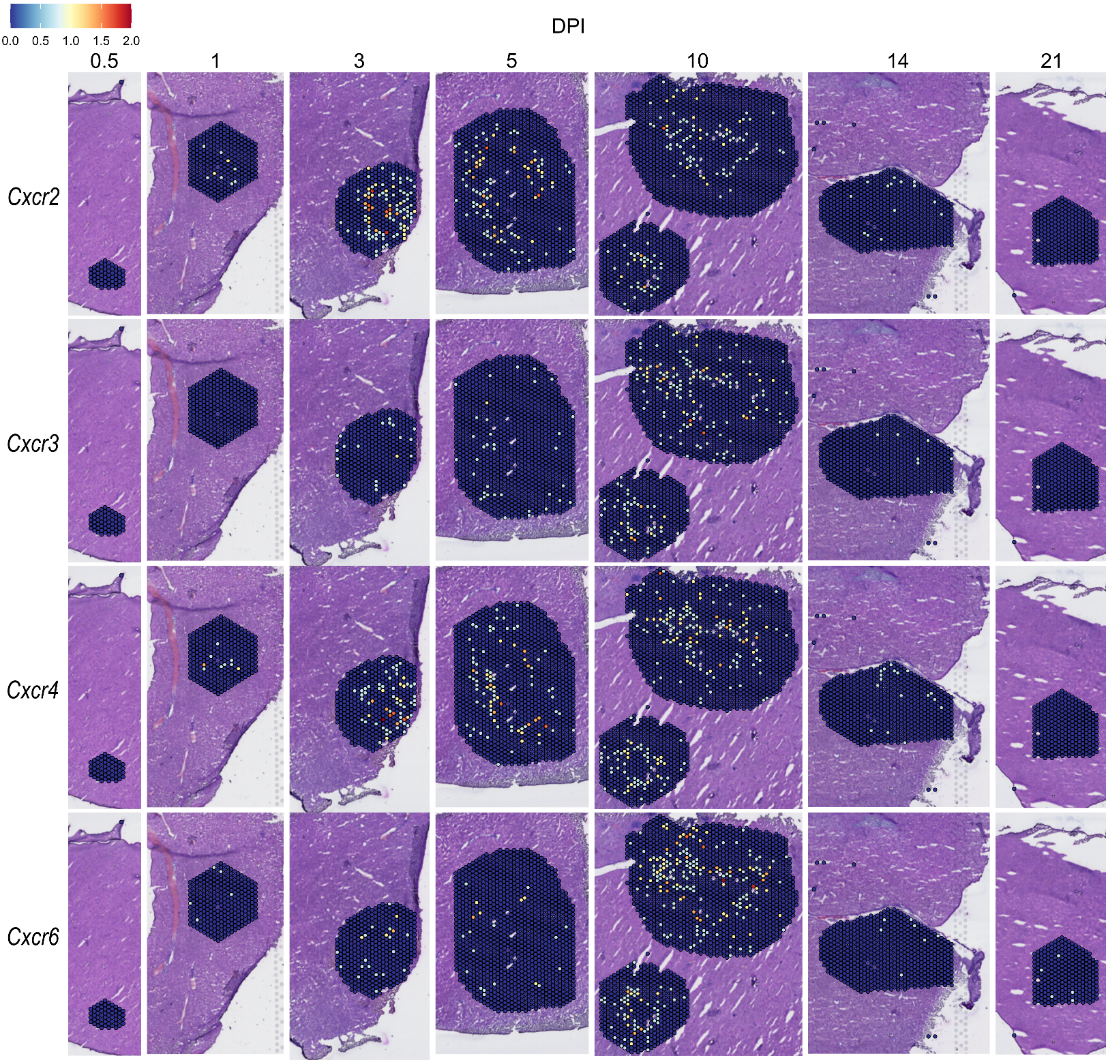

**Figure 4 – figure supplement 5. Spatial expression of *Cxcr* family members.** SpatialFeaturePlots displaying normalized gene expression data of selected *Cxcr* family members at various days post-infection (DPI). Scale set at 0 – 2.0 expression.

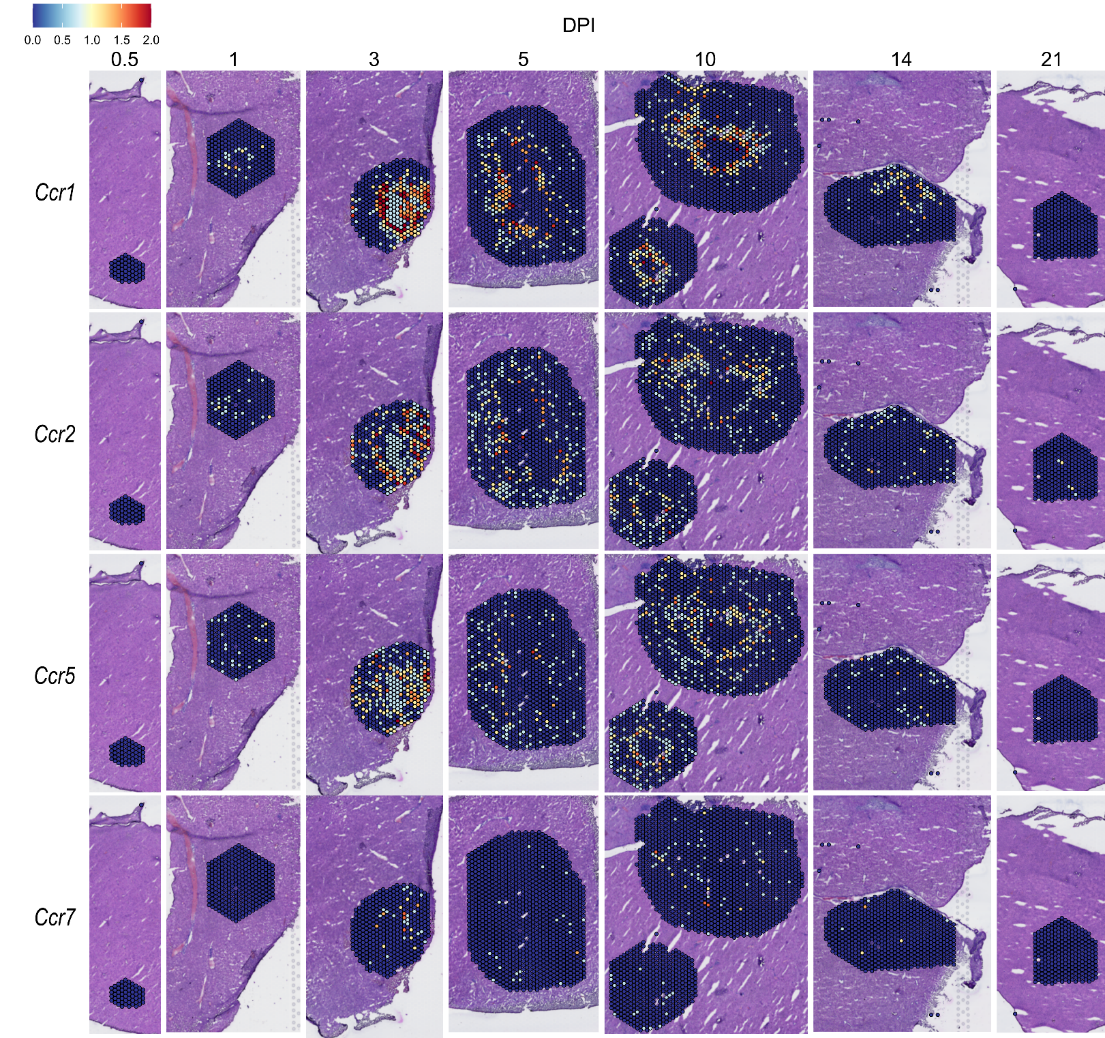

**Figure 4 – figure supplement 6. Spatial expression of *Ccr* family members.** SpatialFeaturePlots displaying normalized gene expression data of selected *Ccr* family members at various days post-infection (DPI). Scale set at 0 – 2.0 expression.

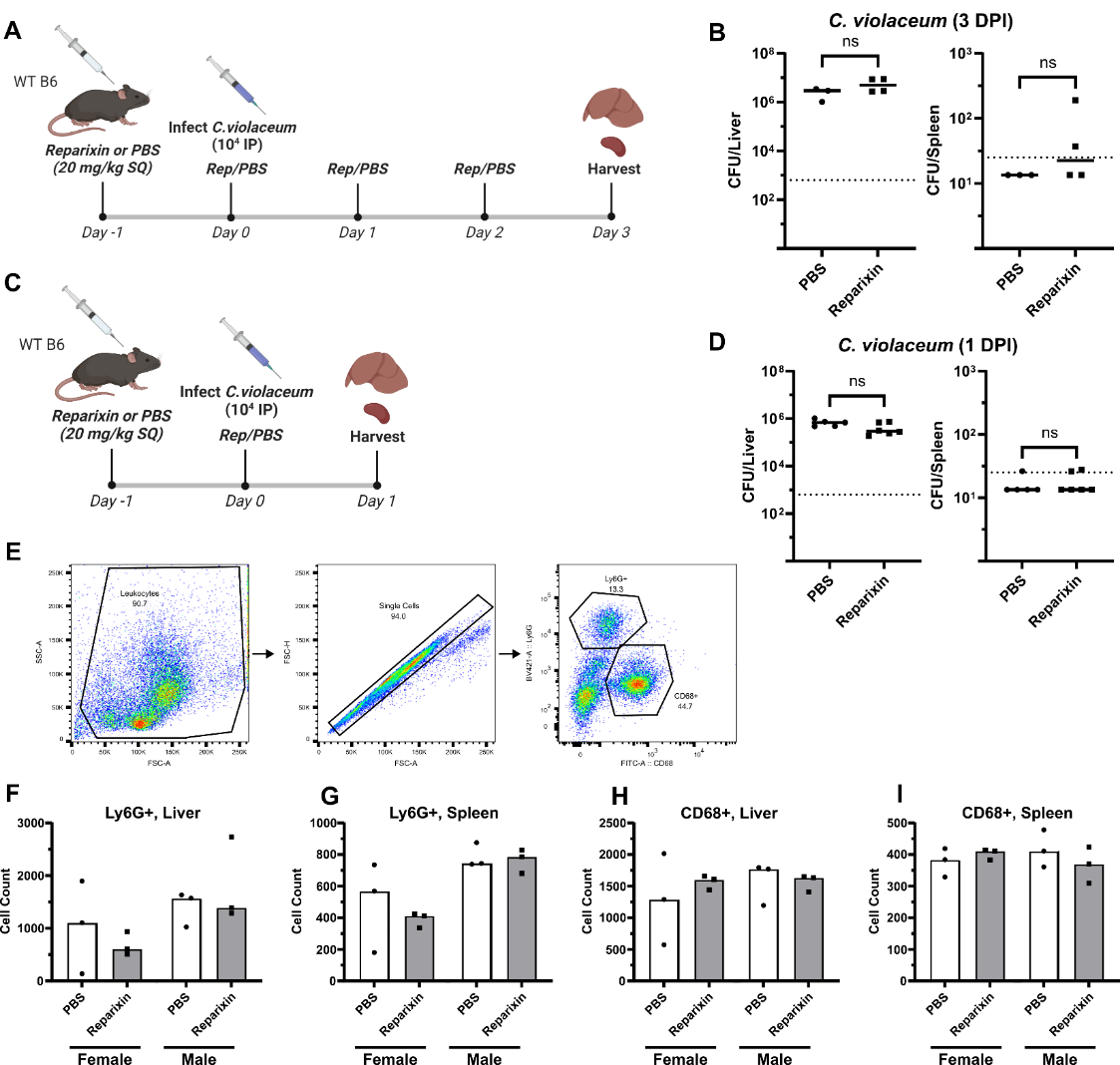

**Figure 5 – figure supplement 1. Reparixin does not inhibit neutrophil chemotaxis into the liver of infected mice.** (**A**) Schematic of the experimental procedure. Mice were injected subcutaneously (SQ) with 20 mg/kg of reparixin, or with PBS. The following day, mice were infected intraperitoneally (IP) with 1x10^4^ CFU of *C. violaceum* and treated again with reparixin or PBS. Mice were treated daily thereafter until harvesting on day 3 post-infection. (**B**) Bacterial burdens in the liver and spleen of PBS- or reparixin-treated mice at 3 days post-infection (DPI). (**C**) Schematic of the experimental procedure as in A, except mice were harvested on day 1 post-infection. (**D**) Bacterial burdens in the liver and spleen of PBS- or reparixin-treated mice at 1 DPI. (**E**) Gating analysis of neutrophil (Ly6G^+^) and macrophage (CD68^+^) numbers via flow cytometry at 1 DPI. Neutrophil numbers in the (**F**) liver and (**G**) spleen. Macrophage numbers in the (**H**) liver and (**I**) spleen. Each dot represents one mouse, with 10,000 events collected per sample. Line at median. (B and D) Dotted line, limit of detection. Solid line, median. Mann-Whitney (abnormally distributed data) for all except liver CFU at 1 DPI, which was analyzed using a two-tailed t test (normally distributed data). Not significant (ns). (B) Liver, p=0.6286; spleen, p=0.4286. (D) Liver, p=0.0641; spleen, p=0.8485. (B) One experiment. (D) Two experiments combined. (F-I) Two experiments combined.

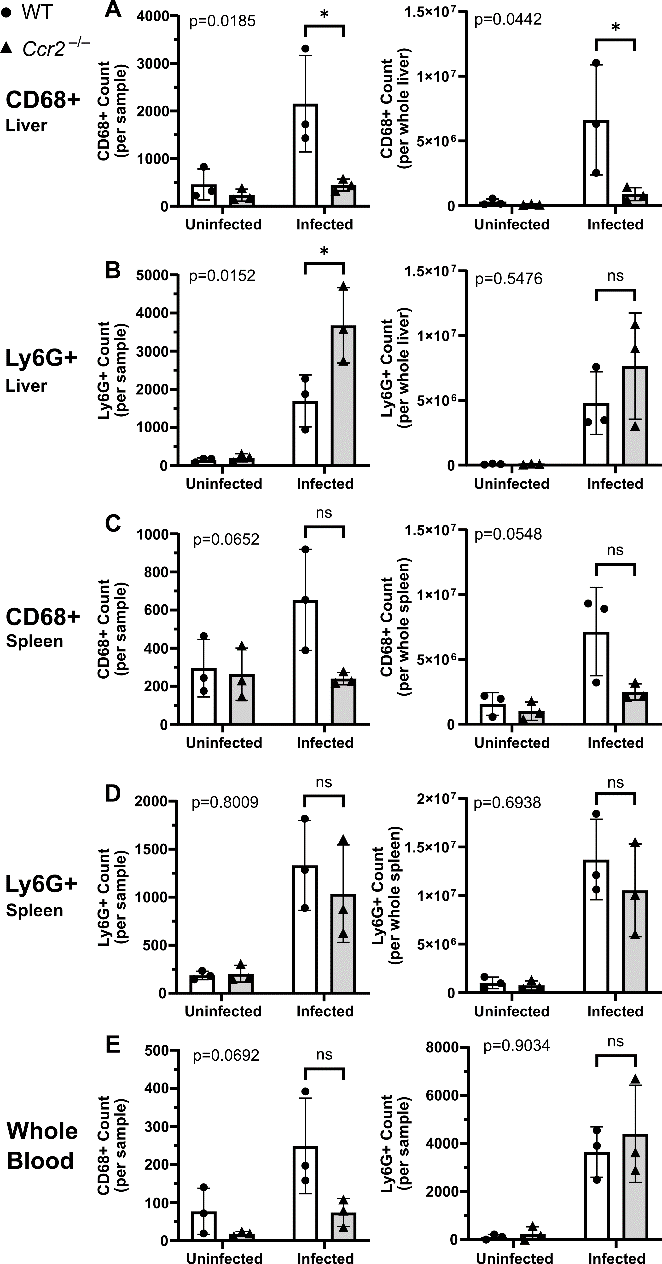

**Figure 6 – figure supplement 1. CCR2 and monocyte recruitment are essential for a successful granuloma response to *C. violaceum*.** Wildtype (WT) and *Ccr2*^–/–^ mice were infected intraperitoneally (IP) with 1x10^4^ CFU *C. violaceum*. (**A-E**) Analysis of macrophage (CD68^+^) and neutrophil (Ly6G^+^) numbers via flow cytometry. Same samples as in Figure 6, but showing cell counts instead of percent; cell counts from single cell gate, with 10,000 events collected per sample, or total cell counts from the whole liver or whole spleen calculated using hemocytometer values following tissue processing; three experiments combined using only female mice, each dot represents one mouse. Two-way ANOVA (for multiple comparisons to assess genotype and infection); key comparisons and p-values shown. Line represents mean ± standard deviation.

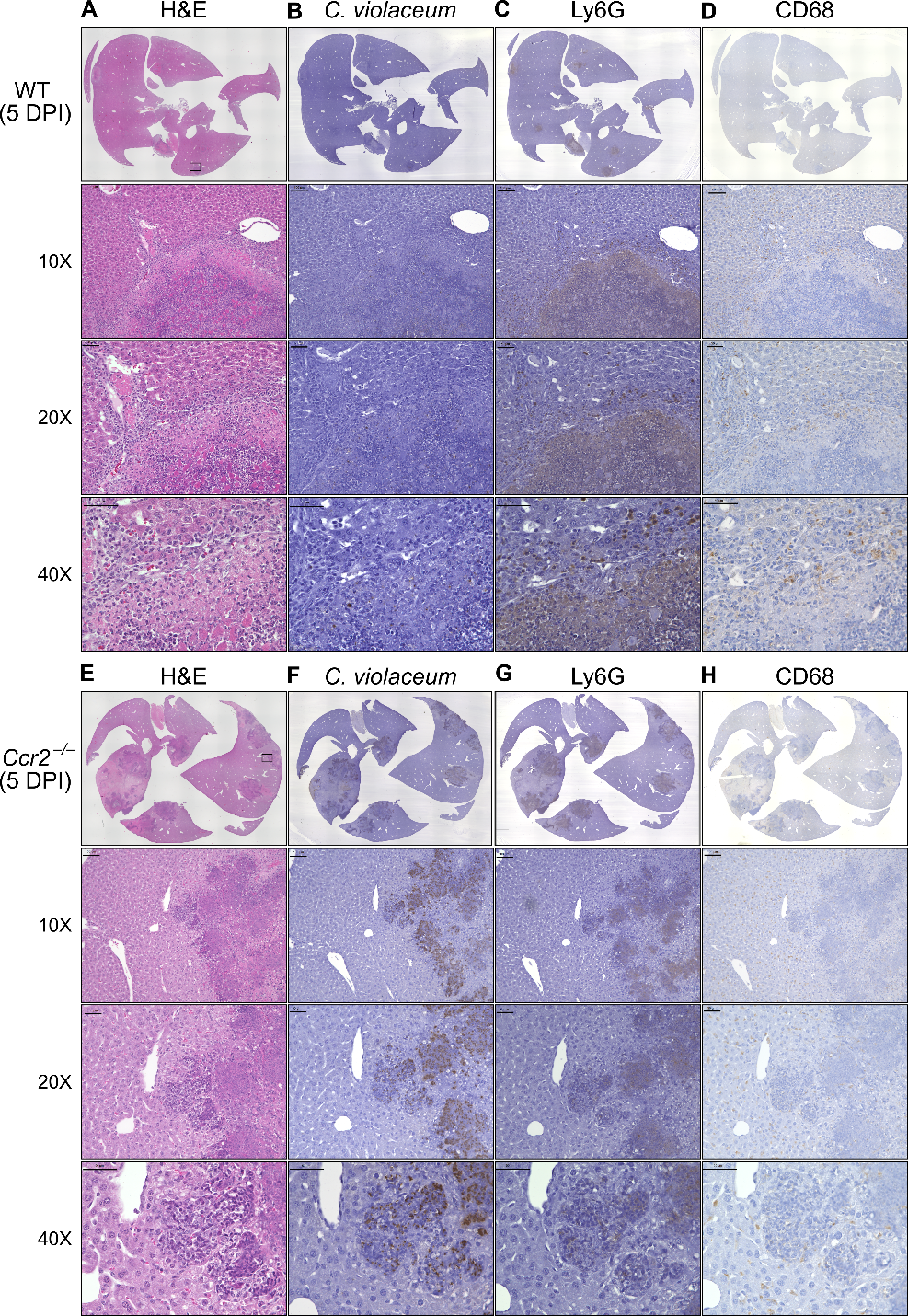

**Figure 7 – figure supplement 1.** **Loss of CCR2-dependent monocyte trafficking results in abnormal granuloma architecture and failure of bacterial containment.** WT and *Ccr2*^–/–^ mice were infected intraperitoneally (IP) with 1x10^4^ CFU *C. violaceum* and livers harvested 5 days post-infection (DPI). Serial sections of livers stained by hematoxylin and eosin (H&E) or various IHC markers for (**A-D**) WT female and (**E-H**) *Ccr2*^–/–^ female. For 10X, scale bar is 100 µm. For 20X and 40X, scale bar is 50 µm.

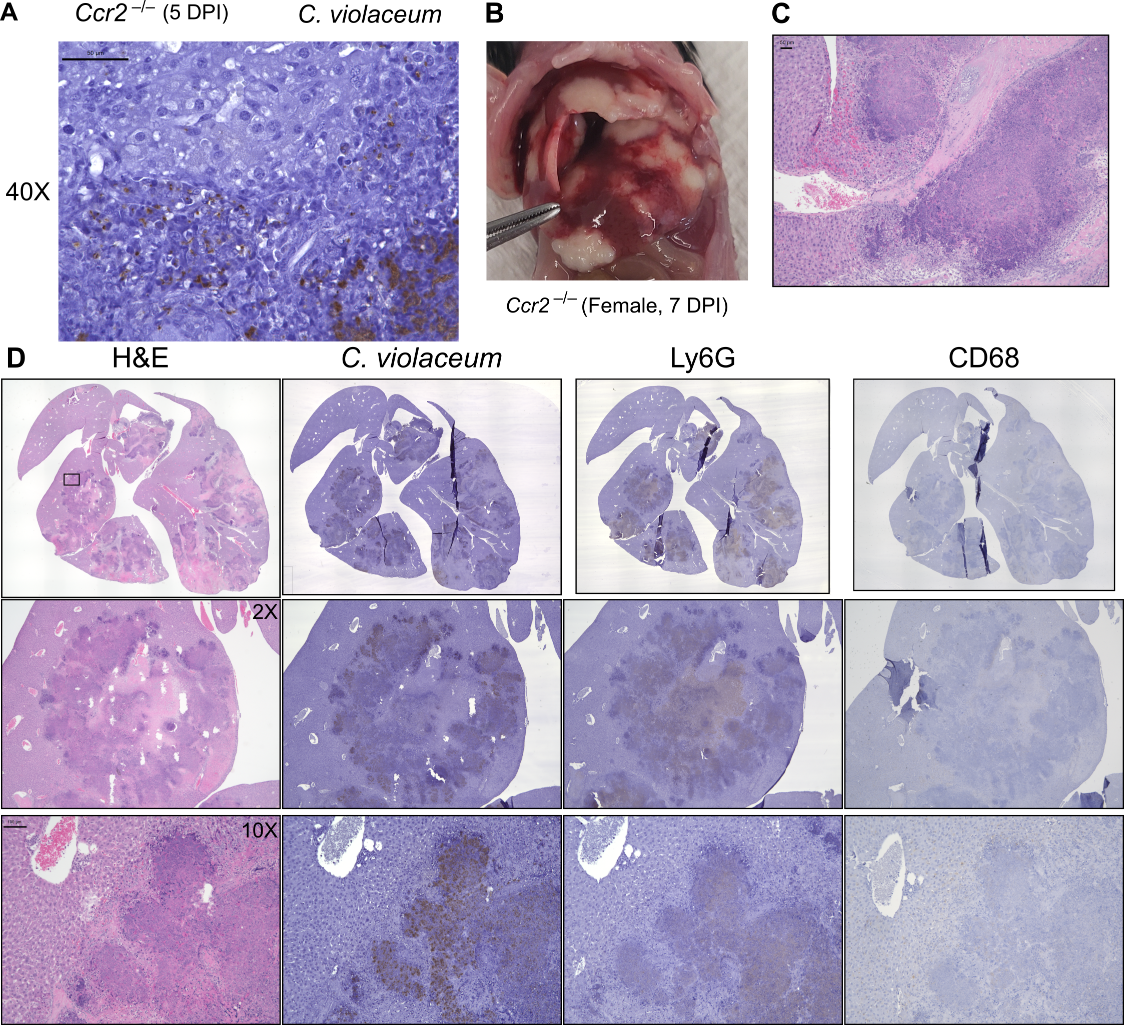

**Figure 7 – figure supplement 2. *Ccr2*^–/–^ mice have increased necrosis and clotting.** *Ccr2*^–/–^ mice were infected intraperitoneally (IP) with 1x10^4^ CFU *C. violaceum*. (**A**) Liver section from *Ccr2*^–/–^ male mouse 5 days post-infection (DPI), stained for *C. violaceum*; zoom showing individual puncta of *C. violaceum*. (**B-D**) A *Ccr2*^–/–^ female mouse from survival curve in Figure 6A that was sacrificed at 7 DPI according to euthanasia criteria. (B) Gross pathology. (C) Liver section stained by hematoxylin and eosin (H&E) showing clotting. (D) Serial sections of liver stained with H&E or various IHC markers. For 10X, scale bar is 100 µm.

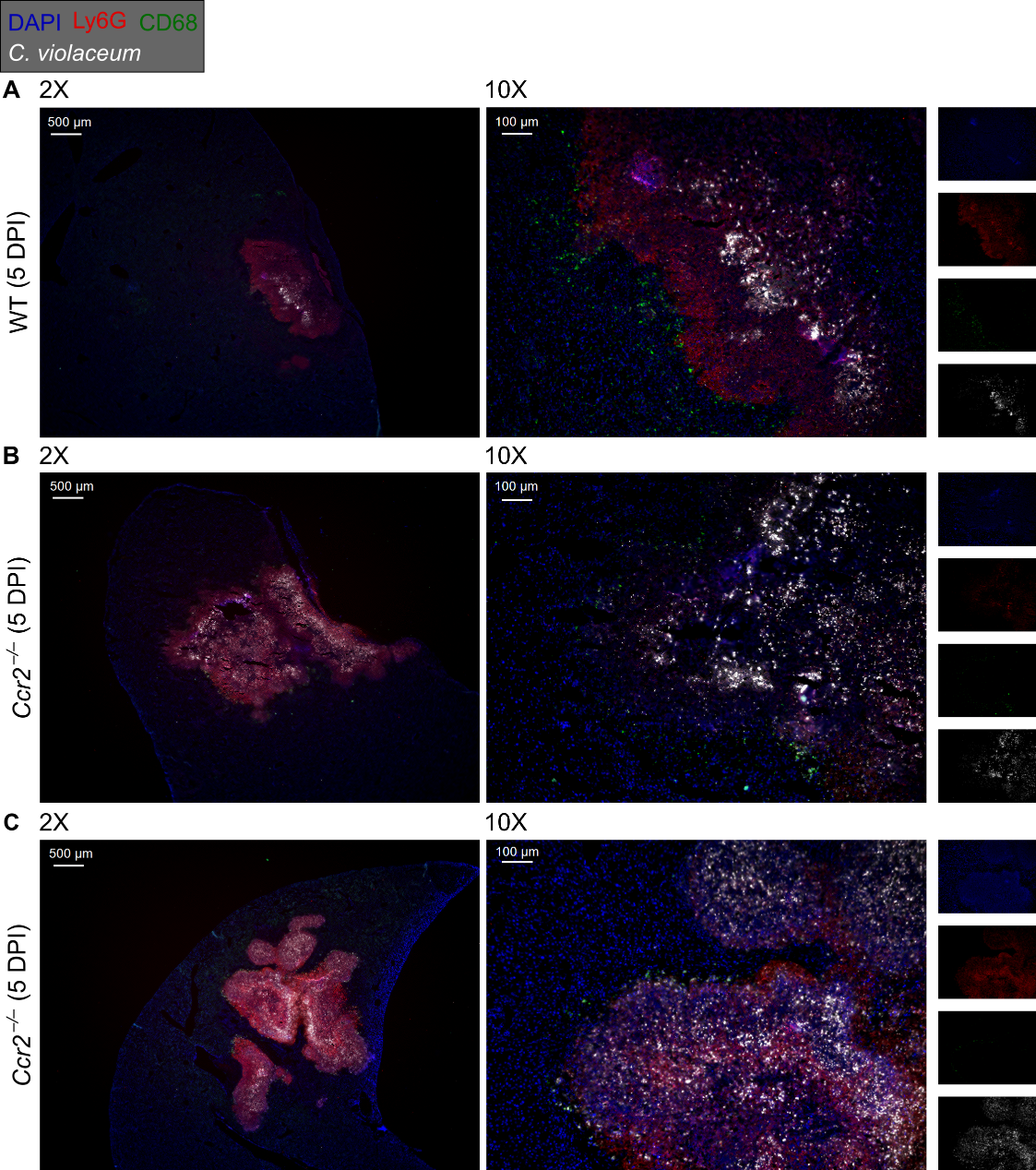

**Figure 7 – figure supplement 3. Loss of CCR2-dependent monocyte trafficking results in abnormal granuloma architecture and failure of bacterial containment.** WT and *Ccr2*^–/–^ mice were infected intraperitoneally (IP) with 1x10^4^ CFU *C. violaceum* and livers harvested 5 days post-infection (DPI) for immunofluorescent staining. Tissue sections were stained for nuclei (DAPI, blue), neutrophils (Ly6G, red), macrophages (CD68, green), and *C. violaceum* (white). (**A**) WT female and (**B-C**) *Ccr2*^–/–^ females. For 2X, scale bar is 500 µm. For 10X, scale bar is 100 µm. Representative of two experiments.

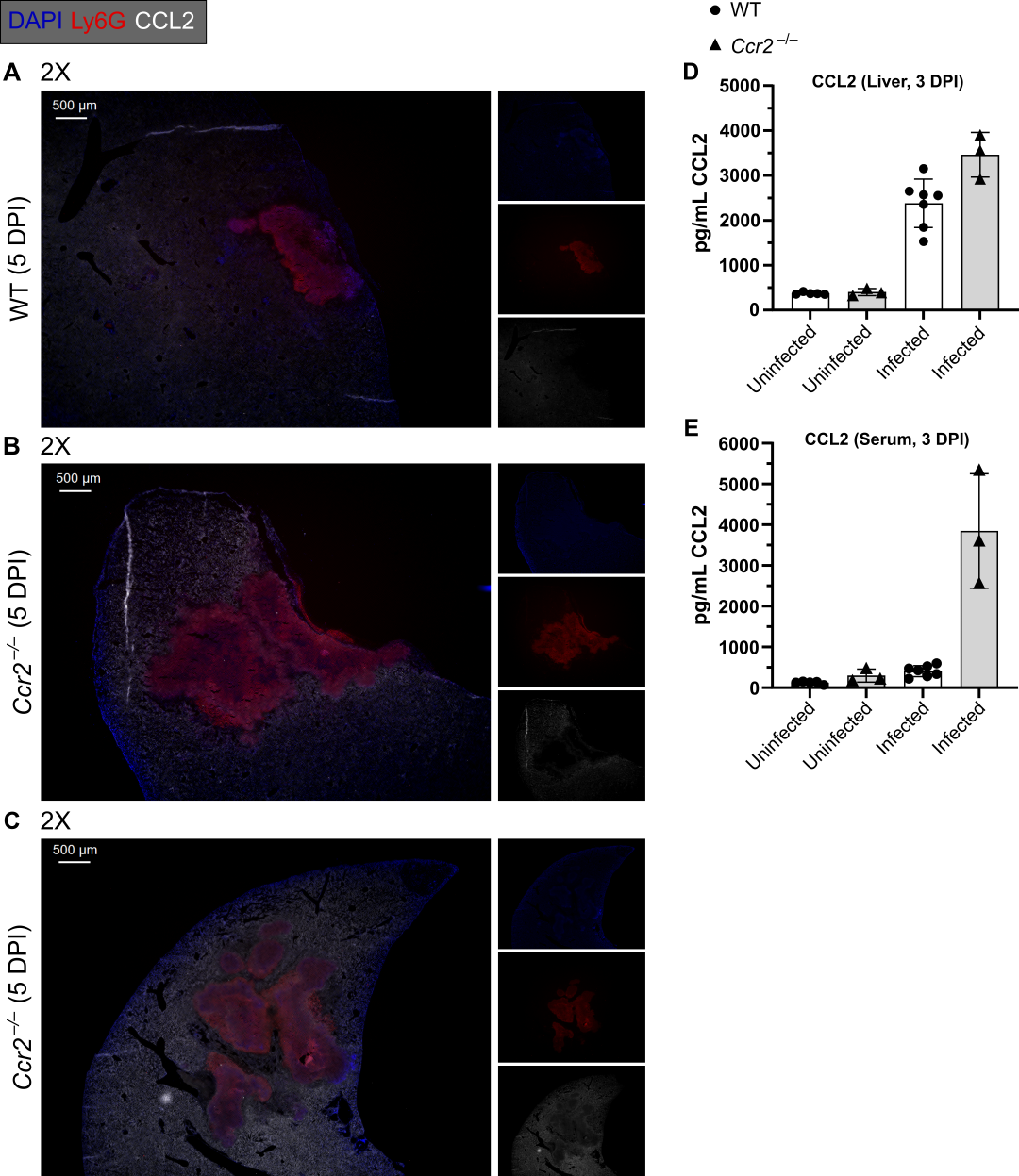

**Figure 7 – figure supplement 4. *Ccr2*^–/–^ mice** **have increased CCL2 in the liver and serum.** WT and *Ccr2*^–/–^ mice were infected intraperitoneally (IP) with 1x10^4^ CFU *C. violaceum*. (**A-C**) Livers harvested 5 days post-infection (DPI) for immunofluorescent staining. Tissue sections were stained for nuclei (DAPI, blue), neutrophils (Ly6G, red), and CCL2 (white). Serial sections from the same tissues in Figure 7 – figure supplement 3. Representative of two experiments. (**D-E**) Quantification of CCL2 via ELISA in the liver (D) and serum (E) of WT and *Ccr2*^–/–^ mice at 3 DPI; two experiments combined using male and female mice, each dot represents one mouse. Line represents mean ± standard deviation.
